# Female and male plumage colour is linked to parental quality, pairing and extra-pair mating in a tropical passerine

**DOI:** 10.1101/2020.06.07.125443

**Authors:** Ana V. Leitão, Michelle L. Hall, Raoul A. Mulder

**Affiliations:** School of BioSciences, University of Melbourne, Parkville, Melbourne, Victoria, Australia; Bush Heritage Australia, Melbourne, Victoria, Australia; School of Biological Sciences, University of Western Australia, Perth, Western Australia, Australia

**Keywords:** Assortative mating, Extra-pair paternity, Female ornaments, Mate choice, Parental Care, Sexual selection

## Abstract

Sexual selection has been proposed to drive the evolution of elaborate phenotypic traits in males, which often confer success in competition or mating. However, in many species both males and females display such traits, although studies reporting selection acting in both sexes are scarce. In this study, we investigated whether plumage ornamentation is sexually selected in female and male lovely fairy-wrens *Malurus amabilis,* a cooperatively breeding songbird. We found that female and male plumage colour was correlated with parental quality but did not incur survival costs. We also found evidence of positive assortative mating based on plumage colour. Microsatellite analyses of paternity indicated that the lovely fairywren has high levels of extra-pair paternity, with 53% of offspring resulting from extra-pair mating. Female and male plumage colour did not predict reproductive success and female proportion of extra-pair offspring in its own nest, but less colourful males obtained higher extra-pair paternity. We argue that plumage colour may be under sex-specific selection, highlighting the importance of looking at both sexes in studies of sexual selection and ornament evolution. The current findings together with previous study, suggest that plumage colour in female and male lovely fairy-wrens appears to be an honest signal relevant in both intra and inter-sexual competition contexts.

## INTRODUCTION

In males of many species, sexual selection is thought to have driven elaborated traits that function in competition or to attract females (Darwin, 1871) by advertising male phenotypic or genetic quality (Andersson, 1994). Although such elaborate traits are more commonly seen in males, they are also expressed in females across many taxa (Amundsen, 2000; Price, 2019). The expression of elaborate traits in females has been often overlooked and thought to be a by-product of selection on males (reviewed in Amundsen, 2000; Amundsen and Pärn, 2006; Tobias et al., 2012). However, more recently, empirical and comparative work suggests that traits are most likely the result of direct selection that can explain female functional signals, either to increase reproductive success (sexual selection; Fitzpatrick and Servedio, 2017; Hare and Simmons, 2019), or to increase access to limited resources via competition (social selection; LeBas, 2006; Tobias et al., 2012; West-Eberhard, 1983).

Morphological traits, such as plumage colours, may be used to reliably assess potential mates if they are correlated with aspects of individual quality (Hunt et al., 1999; Kraaijeveld et al., 2004; MacDougall and Montgomerie, 2003). Males, like females, may benefit from choosing mates based on direct benefits such as phenotypic or genetic quality, which could affect their reproductive success (Amundsen and Pärn, 2006). Traits may also serve as status signals and evolve through competition over mates or other resources (Tobias et al., 2012). Both social competition and mate choice can lead to positive assortative patterns of pairing and mating because individuals with similar traits are likely to pair or mate more frequently than expected by chance (Burley, 1983).

Many socially monogamous bird species engage in extra-pair mating (Brouwer and Griffith, 2019; Griffith et al., 2002). In species that form pair bonds, but sometimes mate with extra-pair individuals, mate choice may involve both choosing a social partner and the genetic parent(s) for offspring (Moller, 1992). In such a case, extra-pair behaviour may be driven by male and/or female mate choice, though most studies have focused only on males traits and how they may affect paternity (Forstmeier et al., 2014; Griffith et al., 2002; Moreno et al., 2015).

The expression of elaborate traits in males and females may have similar or different selective drivers, signalling the same or distinct information. Studying traits using similar procedures in both sexes offers an excellent opportunity to understand the mechanisms for the evolution of elaborate traits, by way of identifying the differences between sexes in selective pressures and investment trade-offs in ornamentation (Dale et al., 2015; Jacobs et al., 2015; Kraaijeveld et al., 2004; Price and Eaton, 2014; Simpson et al., 2020; Soma and Garamszegi, 2015). However, the mechanisms that have shaped ornamental traits in females are still debated (Price, 2019; Riebel et al., 2019), including whether sexual selection can act on females traits (Clutton-Brock, 2009; Hare and Simmons, 2019; Tobias et al., 2012).

Here, we studied the role of plumage colour of female and male lovely fairy-wren, *Malurus amabilis,* a tropical socially monogamous and cooperatively breeding songbird. The lovely fairy-wren is dichromatic, with both male and female displaying conspicuous, but qualitatively different forms of colouration. In both sexes, plumage colouration is correlated with aggression in the context of same-sex competition (Leitão et al., 2019a), but its role in mate choice is unknown. In several closely-related fairy-wren species, the timing of moult into nuptial plumage in males is a sexual signal and a predictor of extra-pair mating success (Brouwer et al., 2011; Cockburn et al., 2008; Dunn and Cockburn, 1999; Karubian, 2002). However, in lovely fairy-wrens, males (and females) retain their bright plumage year-round after reaching maturity, and do not undergo a seasonal moult (Leitão et al., 2019b) so other traits, such as plumage colour, are likely to be implicated as sexual signals. The species’ mating system was unknown before the present study, but fairy-wrens (*Malurus*) exhibit extremely high but variable rates of extra-pair paternity (EPP), ranging across species from 4.4% to 76% of offspring (Cockburn et al., 2013; Kingma et al., 2009; Mulder et al., 1994; Rowe and Pruett-Jones, 2013).

To understand whether plumage colour of both female and male Lovely fairy-wren is a sexual signal, we investigated the links between plumage colour, individual quality, pairing patterns, reproductive success and EPP. Specifically we assessed whether: (i) plumage colour was related to survival and indicators of individual quality; (ii) individuals paired assortatively based on plumage colour; (iii) females and males with the most elaborate plumage had the highest reproductive success; (iv) females mated with extra-pair males on the basis of plumage colour and brighter males had more EPP.

## METHODS

### Study system and general field methods

Lovely fairy-wrens are a non-migratory species endemic to the wet tropics of North Queensland, Australia. Field work was conducted in the Cairns area in Far North Queensland (16.87°S, 145.75°E), where a colour-banded population was established in 2013 and monitored annually until 2017 (for further details see Leitão et al., 2019b).

Lovely fairy-wrens defend territories year-round and breed virtually all year, but with a peak in activity in the Austral Spring, between August and November (Leitão et al., 2019b). Progeny may remain on their natal territory as helpers and assist in subsequent broods. Delayed dispersal is exclusive to males, and females were never seen to help. In our study population, 34.7% of 49 groups included male helpers (Leitão et al., 2019b).

Birds were captured by targeted mist-netting and individually marked with a numbered metal band (Australian Bird and Bat Banding Scheme) and a unique combination of colour bands. At capture, we measured plumage colour (see below) and collected a small blood sample (less than 50 μl) from the brachial vein which was stored in 100% ethanol for later DNA extraction. To assess body size and condition, we weighed individuals (± 0.5 g) and measured tarsus length (± 0.05 mm).

Once individuals were released, we monitored their territories and group members, nesting behaviour and neighbouring territories (density of breeding pairs) weekly. We estimated annual adult survival (presence or absence) by the recovery rate of banded adult breeders (for details see Leitão et al., 2019b).

When possible, blinded methods were used when recording and analysing behavioural data. For some data, it was not possible to record data blindly since it involved focal behavioural observations.

### Nesting and parental care

Lovely fairy-wrens exhibit bi-parental care, with the major contribution of the male being in post-hatch offspring care. Females build the nests and incubate, while males remain near the nest and often engage in courtship feeding. Females lay 2 to 3 eggs and can re-nest several times a year after unsuccessful and successful attempts (Leitão et al., 2019b).

Nests were located using behavioural observations and monitored every 2-3 days. When nestlings were 7 - 9 day-old, we colour-banded and took a small blood sample for paternity analysis. Putative parents were determined by extensive observations of each pair or group, including their attendance at the nest.

Both males and females defend the nest and provision the nestlings and fledglings (Leitão et al., 2019b). Parental care was quantified by recording nestling provisioning as the number of visits by the female, male and helpers from video recordings (GoPro HD Hero2) of active nests. Adults normally resumed feeding within a maximum of 10 min of placing the camera and could be distinguished by their colour bands. We recorded nests for about one hour in the morning (between 0620-1230) when nestlings were 2-5 days (early stage, n=10) and 7-9 days (late stage, n = 9) old. As much as possible, we recorded food provisioning two times per nest (n=6), but some nests do not have repeated measures due to predation of later stage nestlings (n=4), or because the breeding attempt was found at later stages (n=3). For the analysis, we only included nests from pairs that did not have helpers (n = 2 excluded).

### Microsatellite analysis, paternity assignment, and genetic similarity

Genomic DNA was extracted from blood samples using a standard salt extraction method (Bruford et al., 1992). In total, 209 individuals were genetically screened by the Australian Genome Research Facility (Melbourne, Australia) for up to 25 loci (Appendix: Table A1). From these, we selected 14 loci for analysis (Table 1) that had high variability, high call rates, did not deviate significantly from Hardy–Weinberg equilibrium, and had low null allele frequencies (Fnull < 0.03, except for Smm1, Mala14, and Mcymu8 where Fnull = 0.073-0.118; this was taken into account during paternity assignment). For each locus, we calculated the probability of maternal and paternal exclusion, based on allele frequencies of 117 breeders. In total across all 14 loci, we had 173 alleles (X = 12.357) and the combined probability of incorrectly assigning a random male as the sire was less than 1% (Table 1).

**Table 1.** Description of microsatellite loci used for parentage analyses, based on 117 dominant, putatively unrelated birds.

| Locus | N<br>individuals<br>typed | Nr<br>Alleles | Heterozygosity |  | Maternal<br>exclusion<br>probability | Paternal<br>exclusion<br>probability | Null allele<br>frequency |
| --- | --- | --- | --- | --- | --- | --- | --- |
|  |  |  | Expected | Observed |  |  |  |
| Mcymu7<sup>1</sup> | 116 | 4 | 0.573 | 0.543 | 0.163 | 0.282 | 0.023 |
| Mcymu8<sup>1</sup> | 111 | 22 | 0.917 | 0.721 | 0.701 | 0.824 | 0.118 |
| Tmm6<sup>2</sup> | 111 | 58 | 0.971 | 0.946 | 0.875 | 0.933 | 0.011 |
| Smm1<sup>3</sup> | 83 | 7 | 0.621 | 0.518 | 0.225 | 0.404 | 0.079 |
| MaLa2<sup>4</sup> | 115 | 17 | 0.874 | 0.861 | 0.588 | 0.742 | 0.004 |
| MaLa4<sup>4</sup> | 115 | 5 | 0.637 | 0.652 | 0.216 | 0.361 | -0.016 |
| MaLa5<sup>4</sup> | 114 | 6 | 0.63 | 0.658 | 0.207 | 0.354 | -0.021 |
| MaLa6<sup>4</sup> | 115 | 5 | 0.671 | 0.67 | 0.242 | 0.404 | -0.002 |
| MaLa7<sup>4</sup> | 115 | 5 | 0.482 | 0.513 | 0.122 | 0.274 | -0.033 |
| MaLa8<sup>4</sup> | 115 | 8 | 0.699 | 0.73 | 0.277 | 0.444 | -0.025 |
| MaLa10<sup>4</sup> | 115 | 13 | 0.827 | 0.809 | 0.486 | 0.658 | 0.007 |
| MaLa13<sup>4</sup> | 115 | 8 | 0.772 | 0.809 | 0.39 | 0.571 | -0.030 |
| MaLa14<sup>4</sup> | 115 | 9 | 0.808 | 0.704 | 0.452 | 0.629 | 0.073 |
| MaLa18<sup>4</sup> | 114 | 6 | 0.763 | 0.763 | 0.369 | 0.549 | -0.003 |
| Combined | 117 | 173 |  |  | 0.99 | 0.99 | 0.023 |

Maternal exclusion probability is the probability that at a given locus, a random parent will not match the nestling when no parent is known. Paternal exclusion probability is the probability that a candidate father will be excluded assuming that the mother is known. References for microsatellites are: <sup>1</sup>(Double et al., 1997); <sup>2</sup>(Adcock and Mulder, 2002); <sup>3</sup>(Maguire et al., 2006); <sup>4</sup>(Thrasher et al., 2017).

To assign parentage of each offspring (*N* = 58 nestlings) we assumed that the female that built, incubated, and fed the nestlings was the genetic mother (in this species, groups have only one adult female). Offspring were only included in the analysis if they mismatch with the putative mother at 2 or fewer loci. We also examined parentage in groups with juveniles (fledglings or subordinates *N* = 19), to determine if they were likely to be offspring of the breeding pair. For both cases, the social male was accepted as the sire if the nestling mismatched the male’s genotype at 2 or fewer loci. Additionally, to assign potential extrapair sires, we used CERVUS 3.0.7. (Kalinowski et al., 2007; Marshall et al., 1998) to select the most likely male sire from the population, by using the log likelihood score (LOD) for each male in the population and comparing to the offspring (pair LOD score). Each paternity assignment was then checked, and we accepted the assignment in 69 out of 77 cases where the selected male had less than two mismatches. In 4 cases no male was assigned, and in 4 other cases we did not accept the assigned male and in 3 of these we did not assign paternity. Evidence suggested that the assigned male was not the most likely sire, for e.g., sibling from the same brood or future brood (N = 2),a male from a distant population (N = 1; from more than 5 km away, maximum natal and breeder dispersal observed was 1.2 km (Leitão et al., 2019b)), or a male that we knew was dead (N = 1).

Rates of EPP for nestlings and juveniles were analysed separately, but results were similar (Table 3). In total, we were able to assign paternity for 70 out of 77 nestlings and juveniles.

**Table 2.** Parental care (number of visits to the nest) of females and males. Significant predictors in bold, marginal to significance italicised. *N* = 24

| Full Model | $\beta$ | SE | $z$ | $P$ |
| --- | --- | --- | --- | --- |
| Intercept | -7.70 | 0.50 | -15.37 | < **0.001** |
| Sex<sup>a</sup> | 0.08 | 0.39 | 0.21 | 0.83 |
| Cheek colour | 0.27 | 0.10 | 2.62 | **0.008** |
| Head colour | -0.06 | 0.04 | -1.48 | 0.13 |
| Nestling age | 0.10 | 0.05 | 1.93 | *0.053* |
| Brood size | 0.03 | 0.15 | 0.25 | 0.79 |
| Sex<sup>a</sup> * Cheek | -0.63 | 0.48 | -1.30 | 0.19 |
| Sex<sup>a</sup> * Head | 0.02 | 0.09 | 0.23 | 0.81 |
| **Random Effect** | $\sigma^2$ | | | |
| Individual id | 0.00 ($N = 14$) | | | |
| Nest id | 0.00 ($N = 9$) | | | |
| Reduced Model | $\beta$ | SE | $z$ | $P$ |
| Intercept | -7.56 | 0.35 | -21.26 | < **0.001** |
| Sex<sup>a</sup> | 0.05 | 0.39 | 0.15 | 0.87 |
| Cheek colour | 0.28 | 0.10 | 2.73 | **0.006** |
| Head colour | -0.05 | 0.03 | -1.56 | 0.11 |
| Nestling age | 0.09 | 0.05 | 1.90 | *0.057* |
| Sex<sup>a</sup> * Cheek | -0.59 | 0.39 | -1.51 | 0.13 |
| **Random Effect** | $\sigma^2$ | | | |
| Individual id | 0.00 ($N = 14$) | | | |
| Nest id | 0.00 ($N = 9$) | | | |

<sup>a</sup> Sex is categorical term and reference is female

**Table 3.** Frequency of extra-pair nestlings and juveniles (sired by male other than the female’s social partner).

|  | **Nestlings** | **Juveniles** | **Total** |
| --- | --- | --- | --- |
| EPP offspring | 53.44 % (31 out of 58) | 31.57 % (6/19) | 48.05 % (37/77) |
| Broods with at least one EP offspring | 58.62 % (17/29) | 33.33 % (4/12) | 51.21 % (21/41) |
| Total offspring sampled | 58 | 19 | 77 |
| Total broods sampled | 29 | 12 | 41 |
| Nr of incomplete broods | 6 | 2 | 8 |

We estimated relatedness within social pairs to assess genetic similarity between females and their social and extra-pair mates, by calculating the pairwise *r* coefficient following the method of Queller and Goodnight (1989) and using the software Coancestry (v.1.0.1.9; Wang, 2011). Pairs were categorised as incestuous (full sibling pairing and parent-offspring) when pairwise relatedness was within the range of the mean ± 1.5 * SD of known first-order relatives (Brouwer et al., 2017) (N = 17, X ± SD = 0.46 ± 0.12, range: 0.30 – 0.70).

### Reproductive and paternity success

We calculated ‘reproductive success’ of each pair as the number of fledglings per year that survived for more than a month. We also calculated male ‘within-pair paternity’ success as the number of offspring a male sired in his own territory; male ‘extra-pair paternity’ success as the total number of extra-pair offspring he sired in the nests of other males, and ‘total paternity’ success as the sum of a male’s within and extra-pair paternity success. These were annual measures for each territory (we did not have more than one surviving brood of the same individual within a year).

### Measures of colouration and analysis

Spectral reflectance properties of female and male plumage were measured at capture using a spectrometer (Ocean Optics JAZ) with an inbuilt light source (PX-3 Pulsed Xenon), connected to a probe and a machined 45° angle end (UV-VIS fiber-optic reflectance). Before measuring each bird, the spectrophotometer was calibrated relative to a white standard (Ocean Optics WS-2) and a dark reference. Methods are described in Leitão *et al.* (2019a). Briefly, we took five readings of different body parts, and summarised reflectance spectra to describe chromatic variation using psychophysical models of avian vision (Vorobyev and Osorio, 1998; Vorobyev et al., 1998). Visual models reduced each spectrum to a set of three (xyz) chromatic coordinates that define its position in avian visual space, where distances are expressed in JNDs (just noticeable differences), using formulas from Cassey *et al.* (2008) as implemented by Delhey *et al.* (2015). Colour vision in birds is mediated by four types of single cones that are sensitive to very short (VS), short (S), medium (M) and long (L) wavelengths. Axis x represents the relative stimulation of the S cone in relation to the VS cone, y axis represents relative stimulation of the M cone in relation to the VS and S cones, and axis the z represents the relative stimulation of the L cone in relation to the VS, S and M cones.

Within each sex, the colours of different (blue) patches are moderately correlated, and cheek and head were the most colourful patches and with higher visual impact in each sex (Leitão et al., 2019a), so we used these two in the analysis. For each sex, we summarised chromatic variation (xyz coordinates) with a Principal Component Analysis (PCA) using a covariance matrix to maintain the JND units of the original data (Delhey et al., 2015). PCA analysis resulted in one factor (PC1_chroma_) that explained between 73 – 99% of chromatic variance, with low PC1_chroma_ values representing high stimulation of the L relative to the VS+S cones for head and cheek, and high stimulation of M relative to VS + S for only the cheek; so high PC1 values indicated individuals with colours richer in shorter wavelengths (UV/blue, Table A2). We ran PCAs separately for each sex, rather than combining the sexes (cf appendix and main text analyses in Leitão et al., 2019a), because females and males are dichromatic, and we sought to test for patterns of assortative mating between sexes without combining them in a PCA.

In a subsample of re-captures, we found that colour was moderately repeatable between years (Appendix Table A3), so we pooled colour data from different years to generate a single measure per individual.

### Statistical analysis

All statistics were performed in R (version 3.4.4) and Rstudio (version 1.1.414).

To assess whether plumage colouration indicates individual quality or is related with survival costs, we tested associations between plumage colours and multiple response variables representing (1) body condition, estimated by the scaled mass index following (Peig and Green, 2009), (2) body size, (3) parental care, and (4) survival. For the response variables body size and body condition, we built a linear model with normal error distribution for females and males separately and included cheek and head colour as predictors. For the parental care analysis, we built a general linear mixed model (GLMM) with Poisson distribution and individual number of feeding visits as response variable, and included the explanatory variables cheek and head colour and its interaction with sex of the parent, brood size and nestling age. We also included the time of observation as an offset to control for differences in the duration of the observations, and individual ID and nest ID as random effects to account for repeated measures of the same individual and nest. To test for individual survival, we used a general linear mixed model with survival as a binomial response variable (survived more than 12 months after capture/colour measurement = 1; died before one year = 0), cheek colour, head colour and its interaction with sex as predictors, and with year as a random effect, to account for annual differences in survival probability.

We analysed patterns of assortative pairing and mating, by testing Pearson’s correlations between females and their social partner and extra-pair mates in terms of colour (cheek and head plumage), body size (tarsus length) and body condition.

To test for associations between plumage colouration and female and male reproductive success (number of fledglings per year), we used a GLMM with Poisson distribution, to test the predictors plumage colour of both female and male (cheek and head colour) and their interaction, presence of helpers, and territory size (as a measure of territory quality). We included female ID and male ID as random factors to account for repeated measures of the same individuals in different years.

Previous work suggests that there is no single explanation for variation in EPP (Brouwer and Griffith, 2019; Brouwer et al., 2017), so here we considered several possible predictors, while testing whether individual variation in EPP is related to ornament expression. We analysed the proportion of extra-pair offspring in a brood to understand if female plumage colour relates to number of extra-pair fertilizations. We did this by testing the influence of female’s plumage colour, her partner’s colour, interaction between female and partner’s colour, number of neighbouring territories (measure of density and opportunities), presence of helpers, and territory size on proportion of nestlings in brood that were EP offspring. We fitted the model with binomial distribution and included female ID and male ID as random factors.

To determine which factors influenced male paternity success, we used three GLMMs with normal error distribution and log transformed response variables for the total number of offspring that a male sired: ‘within-pair paternity’, ‘extra-pair paternity’ and ‘total paternity’ (within + extra-pair). We assessed the effects on paternity of the male’s own plumage colour, partner’s colour, and the interaction between the two, number of neighbouring territories, presence of helpers, and territory size. We included male ID and partner’s ID as a random factor. Because of the low number of incestuous pairs and general low relatedness between individuals (see Results), we did not include relatedness in the analysis.

In the models of parental care, reproductive success, female mate choice and male paternity success, we present reduced models, due to the higher number of predictors tested, where non-significant terms *(P* > 0.2) were sequentially removed from the models by stepwise backward procedures, starting with interactions terms. We also present full models for comparison.

We compared extra-pair sires with the within-pair male, using a Wilcoxon paired-test, for plumage colour (head and cheek), body size, body condition, genetic similarity to the female, and reproductive success.

GLMs were conducted using packages lme4 (Bates and Maechler, 2010) and the significance of factors and degrees of freedom were assessed using the lmerTest package (Kuznetsova et al., 2017). Differences in sample sizes between tests are due to incomplete data on reflectance spectra, physical measurements, or parental care for some individuals.

## RESULTS

### Plumage colour, individual quality, and survival

We found no relationship between plumage colour and body condition or body size in females and males (Appendix Table A4). Colour of the cheek patch was positively correlated with parental care, with more colourful parents feeding their nestlings more (Figure 1, Table 2). No association was found between survival and plumage colour or its interaction with sex (Appendix Table A5).

**Figure 1.**
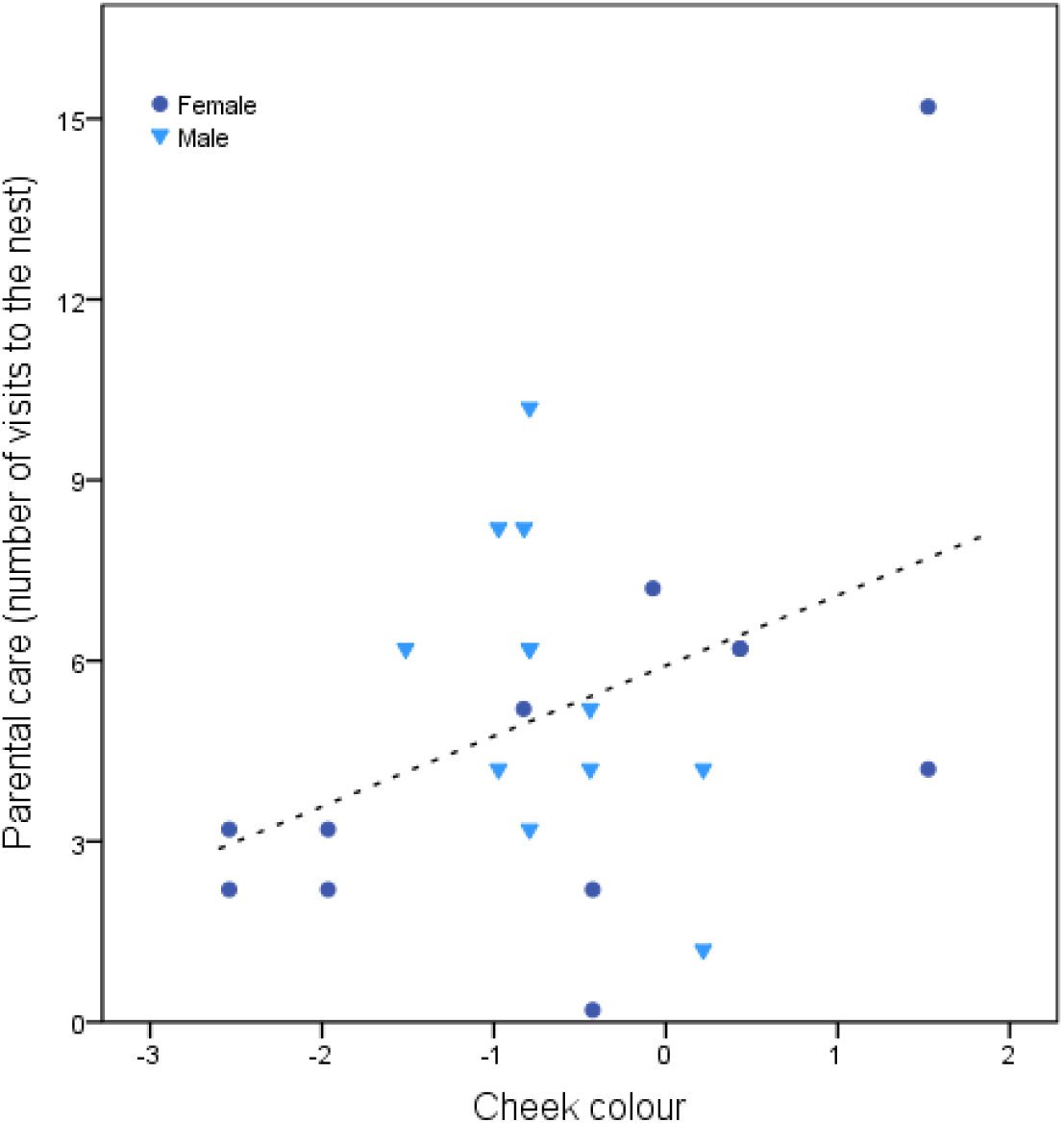
Higher colour of the cheek is related to increase in parental care (number of visits to the nest) of females and males. Dashed lines depict the linear regression line.

### Assortative pairing

Females with more colourful cheek patches paired with males who also had bluer cheeks (female cheek and male cheek: *r* = 0.32, *N* = 36, *P* = 0.05, Figure 2). No relation was found between female and male head colour patch (female head and male head: *r* = 0.07, *N* = 36, *P* = 0.66). Additionally, no patterns of assortative pairing were found based on body size (tarsus length *r* = 0.02, *N* = 37, *P* = 0.91), or body condition (*r* = −0.15, *N* = 37 *P* = 0.36).

**Figure 2.**
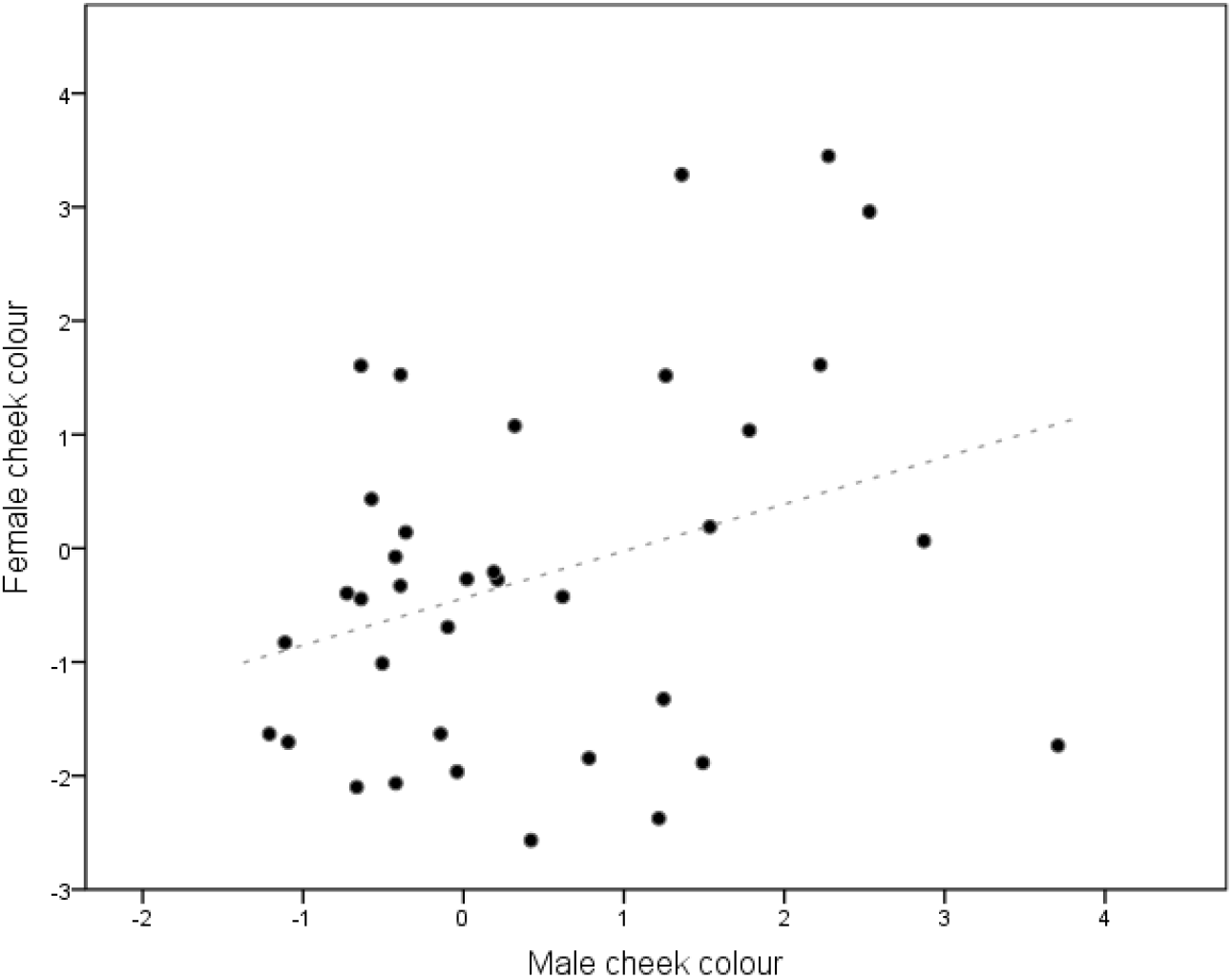
Assortative (positive) pairing based on plumage colour of the female cheek and male cheek. Dashed line shows regression line.

### Frequency of extra-pair paternity and relatedness

Paternity analysis showed that 53% of 58 offspring resulted from extra-pair mating and 58% of 29 broods from 26 groups analysed contained offspring sired by males other than the female’s social partner (Table 3). Of 19 juveniles from 11 groups analysed, 32% were extrapair. Most broods were fathered by a single male (72% of 29 broods: 12 by the social pair male only and 9 by a single extra-pair male). The remaining broods were fathered by up to three different males: 5 broods were fathered by the social male and one extra-pair male; 3 broods were fathered by multiple extra-pair males.

Ten percent of pairs were incestuous pairings (3 of 33 pairs where both male and female were genotyped). Five out of 7 nestlings in the 3 broods from these pairs were EP offspring.

### Reproductive and paternity success

The number of surviving offspring produced by the pair was unrelated to female or male plumage colour or any other variable tested (Table 4). Females had more extra-pair offspring when they had more neighbours (Table 5), and the proportion of nestlings in the brood that were EP offspring did not depend on the female’s or her mate’s colour or any other variable tested, yet in the reduced model less colourful females tended to have more EPP offspring (Table 5).

**Table 4.** Annual reproductive success of females and males (number of fledglings produced that survived). Significant predictors in bold, marginal to significance italicised. *N* = 26.

| **Full Model** | $\beta$ | SE | $z$ | $P$ |
| --- | --- | --- | --- | --- |
| Intercept | -1.66 | 1.40 | -1.18 | 0.23 |
| Female cheek colour | 0.25 | 0.19 | 1.31 | 0.19 |
| Female head colour | -0.39 | 0.44 | -0.88 | 0.37 |
| Male cheek colour | -0.48 | 0.31 | -1.52 | 0.12 |
| Male head colour | -1.35 | 0.83 | -1.61 | 0.10 |
| Presence of helpers<sup>a</sup> | 0.55 | 0.52 | 1.06 | 0.28 |
| Territory size | 0.11 | 0.12 | 0.92 | 0.35 |
| Female cheek * Male cheek | 0.19 | 0.23 | 0.83 | 0.40 |
| Female head * Male head | -0.41 | 0.25 | -1.62 | 0.10 |
| **Random Effect** | $\sigma^2$ | | | |
| Female ID ($N = 16$) | 0.00 | | | |
| Male ID ($N = 19$) | 0.00 | | | |
| **Reduced Model** | $\beta$ | SE | $z$ | $P$ |
| Intercept | 0.70 | 0.35 | 1.97 | **0.04** |
| Female head colour | -0.44 | 0.36 | -1.22 | 0.22 |
| Male head colour | -1.13 | 0.69 | -1.63 | 0.10 |
| Female head * Male head | -0.36 | 0.20 | -1.74 | *0.08* |
| **Random Effect** | $\sigma^2$ | | | |
| Female ID | 0.00 |  |  |  |
| Male ID | 0.00 |  |  |  |

<sup>a</sup> Helper is categorical term and reference is absence of helpers

**Table 5.**
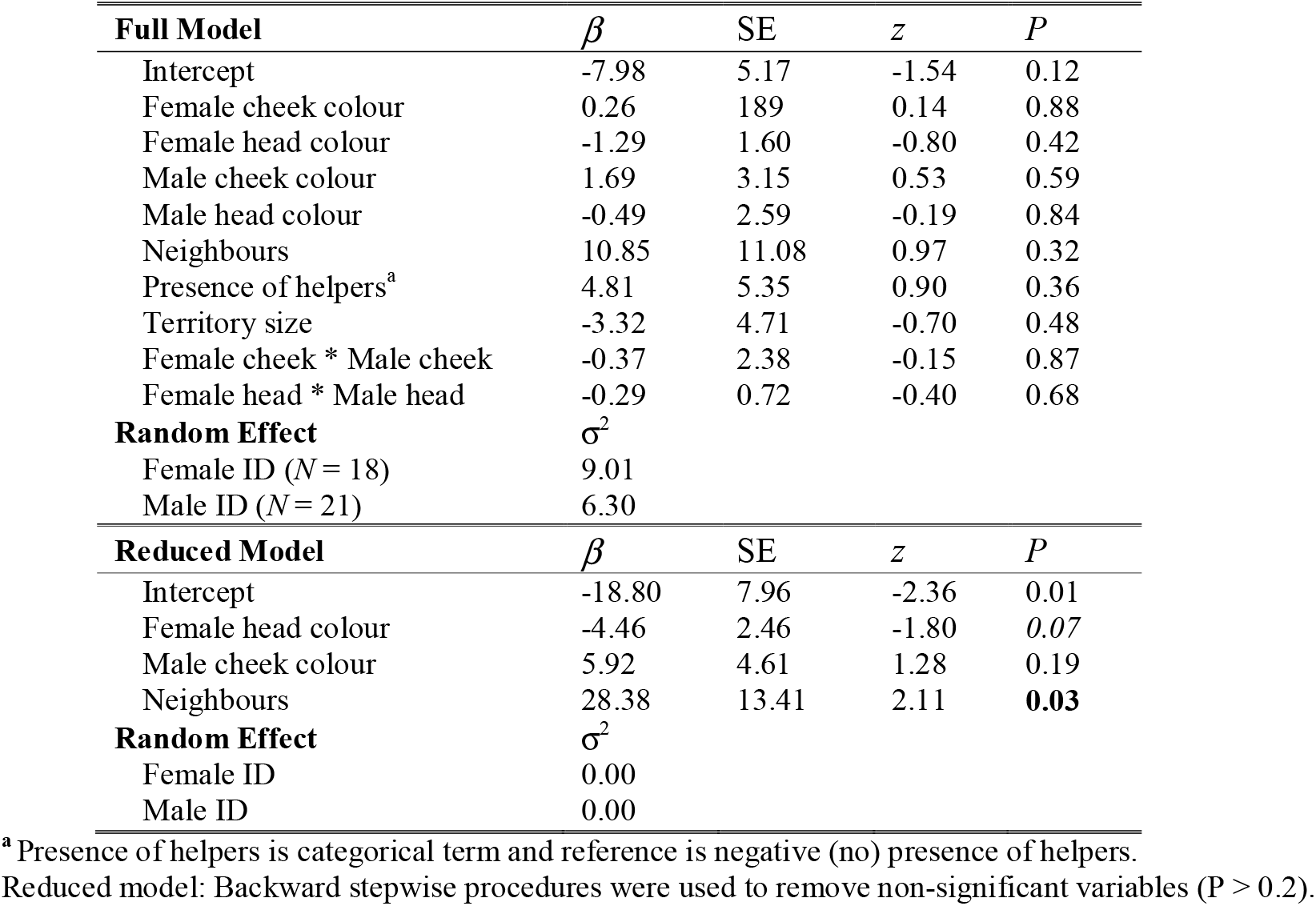
Proportion of nestlings in brood that were EP offspring. Significant predictors in bold, marginal to significance italicised. *N* = 28.

| **Full Model** | $\beta$ | SE | $z$ | $P$ |
| --- | --- | --- | --- | --- |
| Intercept | -7.98 | 5.17 | -1.54 | 0.12 |
| Female cheek colour | 0.26 | 1.89 | 0.14 | 0.88 |
| Female head colour | -1.29 | 1.60 | -0.80 | 0.42 |
| Male cheek colour | 1.69 | 3.15 | 0.53 | 0.59 |
| Male head colour | -0.49 | 2.59 | -0.19 | 0.84 |
| Neighbours | 10.85 | 11.08 | 0.97 | 0.32 |
| Presence of helpers<sup>a</sup> | 4.81 | 5.35 | 0.90 | 0.36 |
| Territory size | -3.32 | 4.71 | -0.70 | 0.48 |
| Female cheek * Male cheek | -0.37 | 2.38 | -0.15 | 0.87 |
| Female head * Male head | -0.29 | 0.72 | -0.40 | 0.68 |
| **Random Effect** | $\sigma^2$ | | | |
| Female ID ($N = 18$) | 9.01 | | | |
| Male ID ($N = 21$) | 6.30 | | | |
| **Reduced Model** | $\beta$ | SE | $z$ | $P$ |
| Intercept | -18.80 | 7.96 | -2.36 | 0.01 |
| Female head colour | -4.46 | 2.46 | -1.80 | 0.07 |
| Male cheek colour | 5.92 | 4.61 | 1.28 | 0.19 |
| Neighbours | 28.38 | 13.41 | 2.11 | **0.03** |
| **Random Effect** | $\sigma^2$ | | | |
| Female ID | 0.00 |  |  |  |
| Male ID | 0.00 |  |  |  |

<sup>a</sup> Presence of helpers is categorical term and reference is negative (no) presence of helpers.

Males had higher rates of within-pair paternity in less dense areas (number of neighbouring territories), and when groups had no helpers (Table 6). Males had higher EPP when they were in more dense areas, when they had less colourful cheeks, and when assorted negatively with their partner (negative relation between male’s colour and female’s colour, Table 6). Within and extra-pair paternity was negatively correlated, although not significant (Pearson’s *r* = −0.307, *N* = 28, *P* = 0.11). Male total paternity success (within and extra-pair paternity) averaged 1.5 ±1.2 offspring per year (range 0-5, *N* = 28 males) and was marginally negatively related with male cheek colour (Table 6), but not related with any other variable tested (Table 6).

**Table 6.** General linear mixed model showing the predictors of within-, extra-pair and total paternity success of males. Significant predictors in bold, marginal to significance italicised.

|  | Within-pair paternity (*N* = 28 male-years) |  |  |  | Extra-pair paternity (*N* = 30) |  |  |  | Total = Within and extra-pair paternity (*N* = 28) |  |  |  |
| --- | --- | --- | --- | --- | --- | --- | --- | --- | --- | --- | --- | --- |
| **Full Model** | $\beta$ | SE | *t* | *P* | $\beta$ | SE | *t* | *P* | $\beta$ | SE | *t* | *P* |
| Intercept | 0.32 | 0.07 | 4.54 | **<0.001** | 0.10 | 0.06 | 1.58 | 0.13 | 0.40 | 0.07 | 5.10 | **<0.001** |
| Male cheek colour | -0.04 | 0.05 | -0.71 | 0.48 | -0.06 | 0.05 | -1.13 | 0.27 | -0.07 | 0.06 | -1.15 | 0.26 |
| Male head colour | 0.01 | 0.04 | 0.36 | 0.72 | -0.03 | 0.04 | -0.73 | 0.47 | -0.02 | 0.05 | -0.42 | 0.67 |
| Partner's cheek colour | -0.01 | 0.03 | -0.29 | 0.77 | 0.04 | 0.03 | 1.27 | 0.24 | 0.02 | 0.04 | 0.56 | 0.58 |
| Partner's head colour | 0.003 | 0.02 | 0.14 | 0.88 | -0.002 | 0.02 | -0.08 | 0.93 | -0.002 | 0.02 | -0.11 | 0.90 |
| Neighbours | -0.13 | 0.09 | -1.48 | 0.20 | 0.14 | 0.09 | 1.58 | 0.15 | 0.004 | 0.09 | 0.04 | 0.96 |
| Presence of helpers<sup>a</sup> | -0.12 | 0.10 | -1.11 | 0.28 | -0.03 | 0.10 | -0.34 | 0.73 | -0.14 | 0.12 | -1.18 | 0.25 |
| Territory size | -0.001 | 0.09 | -0.02 | 0.98 | 0.06 | 0.08 | 0.79 | 0.44 | 0.01 | 0.09 | 0.14 | 0.88 |
| Male cheek * Partner's cheek | 0.018 | 0.04 | 0.43 | 0.67 | -0.06 | 0.03 | -1.76 | *0.09* | -0.04 | 0.04 | -0.93 | 0.36 |
| Male head * Partner's head | 0.0001 | 0.01 | 0.009 | 0.99 | -0.003 | 0.01 | -0.22 | 0.82 | -0.005 | 0.01 | -0.32 | 0.74 |
| **Random effect** | $\sigma^2$ | | | | $\sigma^2$ | | | | $\sigma^2$ | | | |
| Male ID (*N*) | 0 (21) |  |  |  | 9.69e-11 (22) |  |  |  | 0.0 (21) |  |  |  |
| Female ID (*N*) | 0.008 (18) |  |  |  | 0.009 (18) |  |  |  | 0.0 (18) |  |  |  |
| Residual | 0.03 |  |  |  | 0.02 |  |  |  | 0.05 |  |  |  |
| **Reduced Model** |  |  |  |  |  |  |  |  |  |  |  |  |
| Intercept | 0.32 | 0.04 | 7.82 | **<0.001** | 0.13 | 0.03 | 3.79 | **0.005** | 0.37 | 0.04 | 7.74 | **<0.001** |
| Male cheek colour |  |  |  |  | -0.07 | 0.03 | -2.17 | **0.04** | -0.07 | 0.04 | -1.76 | *0.08* |
| Partner's cheek colour |  |  |  |  | 0.04 | 0.02 | 1.80 | 0.12 |  |  |  |  |
| Neighbours | -0.15 | 0.06 | -2.26 | **0.03** | 0.15 | 0.07 | 2.25 | **0.04** |  |  |  |  |
| Presence of helpers<sup>a</sup> | -0.16 | 0.07 | -2.31 | **0.02** |  |  |  |  | -0.11 | 0.08 |  | 0.19 |
| Male cheek * Partner's cheek |  |  |  |  | -0.07 | 0.02 | -2.54 | **0.02** |  |  |  |  |
| **Random effect** | $\sigma^2$ | | | | $\sigma^2$ | | | | $\sigma^2$ | | | |
| Male ID (*N*) | 0.0 (21) |  |  |  | 0.0 (22) |  |  |  | 0.0 (21) |  |  |  |
| Female ID (*N*) | 0.0 (18) |  |  |  | 0.001 (18) |  |  |  | 0.0 (18) |  |  |  |
| Residual | 0.03 |  |  |  | 0.02 |  |  |  | 0.04 |  |  |  |

<sup>a</sup> Presence of helpers is categorical term and reference is positive (yes) presence of helpers
Reduced model: Backward stepwise procedures were used to remove non-significant variables (*P* > 0.2).

### Extra-pair male traits and comparison with social male traits

Females and their extra-pair males did not mate assortatively based on colour (head - colour: *r* = −0.16, *N* = 12, *P* = 0.60; cheek colour: *r* = −0.01, *N* = 12, *P* = 0.96), or body condition (*r* = −0.22, *N* = 12 *P* = 0.47). Females and extra-pair males tended to mate negatively assortatively by body size (tarsus length: *r* =-0.52, *N* = 11, *P* = 0.09).

A paired comparison between extra-pair males and the female’s social partner showed that extra-pair males tended to be bigger than social partners (respectively 21.63 ± 0.30 and 21.40 ± 0.61 mm tarsus length, *Z=* −1.39, *N* = 16, *P* = 0.08) and have less colourful heads (respectively −1.53 ± 0.99 and −2.50 ± 1.68 head colour, *Z* = −1.47, *N* = 12, *P* = 0.07), but did not differ in cheek colour (*Z* = −1.00, *N* = 12, *P* = 0.15) or body condition (scaled mass index: *Z* = −1.23, *N* = 16, *P* = 0.10). There was also no difference between extra-pair males and social males in their genetic similarity to the female (*Z* = 1.63, *N* = 14, *P* = 0.94).

## DISCUSSION

We investigated whether plumage colouration is a sexually selected signal in female and male Lovely fairy-wrens. We show that more colourful males and females cared more for their offspring, and that females and their social males paired assortatively by plumage colour. We also found high rates of extra-pair paternity, with 53% of nestlings sired by extra-pair males, and that the number of neighbouring territories, but not plumage colour, predicted the number of female’s EP offspring. Males gained more paternity in their own nests at low breeding densities and when they had no helpers in their group. We found that males had higher EPP when they were in more dense areas, and unexpectedly, when they were less colourful. Taken together, our findings suggest that plumage colour signals parental quality and is important in mate choice. Furthermore, males, but not females, may have underlying differences in mating strategies correlated with plumage colouration.

### Plumage colour as positive indicator of parental quality that does not incur survival costs

Female and male ornament expression may be used for mate choice, if it provides information about aspects of phenotypic quality or fitness (Jawor et al., 2004; Laczi et al., 2013). Here, we found that plumage colour predicted parental care, where males and females with more colourful cheek plumages provisioned more. These results, consistent with other studies (Dakin et al., 2016; Linville et al., 1998; Siefferman and Hill, 2003), suggests that plumage colour displayed by both sex is a reliable indicator of parental quality and that individuals can benefit by choosing a mate based on this trait.

We found no further evidence that plumage colour signals size or condition, but it is possible that plumage colouration signals other aspects of quality that were not measured here, such as good genes (Hamilton and Zuk, 1982), immunological capacity (McGraw and Ardia, 2003), or age (Dias et al., 2016). These qualities can co-vary with plumage colour, in such way that individuals are choosing compatible or high-quality sexual partners.

Conspicuous colours are often associated with high survival costs, with several studies suggesting that bright colours attract more predation (Götmark and Hohlfält, 1995; Götmark et al., 1997; Huhta et al., 2003). Still, we found that annual survival was unrelated with plumage colouration. In line with this, an experimental study investigating colour phenotypes in fairy-wrens showed that being brightly coloured did not increase the chances of adult predation (Cain et al., 2019), indicating that increased conspicuousness is not associated with greater predation pressure. The absence of the seasonal plumages characterising other (male) fairywren species in lovely fairy-wrens, paired with living in the tropics and high adult survival (Leitão et al., 2019b), suggests that colourful plumage in the lovely fairy-wren does not incur survival costs.

### Positive assortative pairing: Mate choice for plumage colour and parental qualities?

Consistent with the growing body of evidence that individuals paired assortatively with respect to plumage colouration (de Zwaan et al., 2019; Griggio et al., 2005; Jacobs et al., 2015; MacDougall and Montgomerie, 2003; Rowe and Weatherhead, 2011), we found a similar pattern in the lovely fairy-wrens. Socially monogamous species that contribute similarly to parental care, as seen in this species (Leitão et al., 2019b), are expected to choose a high quality partner (Fitzpatrick et al., 1995; Safran and McGraw, 2004) and in positive assortative mating, individuals of similar traits are expected to pair more frequently than expected by chance (Burley, 1983). When only one sex is selective, patterns of assortative mating are not expected (Burley, 1983). This mating pattern suggests that plumage colour is expressed in both sexes because it has a signalling function and is favoured by mutual mate choice (Kraaijeveld et al., 2007), results that are supported by different empirical work (Heinig et al., 2014; Hunt et al., 1999; Kraaijeveld et al., 2004; MacDougall and Montgomerie, 2003). Yet, the possibility that assortative pairing may be due to reasons other than mutual mate choice cannot be discarded. For example, competition can cause a similar pattern if social dominance affects pair formation (Creighton, 2001). We have shown that plumage colouration is involved in male and female competitive interactions (Leitão et al., 2019a), and thus it can reflect the ability of individuals to compete for territories (Hasegawa et al., 2014; Wolfenbarger, 1999), and territory quality may co-vary with plumage colour. Behavioural observations and experiments are needed to distinguish these possibilities.

### Reproductive success, extra-pair behaviour and correlates with plumage colour

Males with more conspicuous ornaments are assumed to be of better quality because ornaments may be costly to produce (Hill, 2006). Showy males can be more successful when competing for sexual resources and may be preferred as mates (Andersson, 1994; Darwin, 1871), and so it is expected that these individuals achieve higher reproductive success. However, we found that male and female plumage colouration was not correlated with reproductive success in the home nest. This was despite finding a relationship between parental quality and plumage colour, and positive assortative mating by plumage colour. We cannot rule out that our measure of breeding performance did not capture a possible relation or effect with plumage colour. For example, individuals that are brighter could have better quality offspring, but may attract and increase predation at the nest (Wallace, 1889), and so, general reproductive performance may be masked by predation, the main cause of nest failure in this species (Leitão, Hall, Venables, et al., 2019).

If extra-pair males exert a choice, they may favour females that have brighter colours, a preference that can be reflected in higher number of female EP fertilizations. We found that females with more EP offspring in a nest were in areas with high density of males, but no relation was found with female’s plumage colour. These results suggest that males might not be selective or discriminate between different female plumage colours.

Interestingly, we found that less colourful males had higher EPP success. Under female mate choice and assuming that brighter males are of higher quality, it is expected that females prefer less colourful extra-pair males because the benefits of mating with this type of male outweigh the costs (Qvarnström and Forsgren, 1998), for e.g. maintenance costs and higher vulnerability to predators (handicap signals; Zahavi, 1975) or production costs (physiological investment or nutrients) (Doucet, 2002; Keyser and Hill, 2000; McGraw et al., 2002). Another possibility could be that females do not gain much in being selective about extra-pair partners, and EPP is under male control (although in superb fairy-wrens *Malurus cyaneus* EP behaviour is under female control (Double and Cockburn, 2000)). If that is the case, males that are less colourful may be more aggressive, as we have shown is a previous study (Leitão et al., 2019a) and intrinsically willing to engage in extra-pair behaviour more often (Forstmeier, 2007) or obtain copulations by force (Burg and Croxall, 2006).

A male’s reproductive fitness can be determined by the ability to sire and care for their offspring and/or by increasing mating effort (extra-pair fertilisation) (Balshine et al., 2002). Since more colourful males provide more parental care, and less colourful ones have higher extra-pair mating success (negatively correlated), we argue that males may be adopting different reproductive strategies that are correlated to their plumage colour. In such context, different strategies related with male colouration may lead to similar payoffs and comparable final fitness, i.e. number of fledgling success. Since this study was observational, we cannot rule out any of these possibilities, therefore further detailed experimental studies will be needed to test for these. Regardless of the mechanisms, our results suggest that plumage colour is relevant in mate choice in this species, in both social and extra-pair contexts.

### Lovely fairy-wren mating system

We found that lovely fairy-wrens had high rates of EPP despite low frequencies of extra-pair courtship displays, high male care, and low number of helpers compared to other *Malurus* (Leitão et al., 2019b). Previous comparative work in the Maluridae family (which did not include the lovely fairy-wren) showed that multiple factors predicted the variation of EPP rates, including inbreeding avoidance, the number of helpers, male care, and male density (Brouwer et al., 2017). We found similar predictors in this study, since males that were in areas with lower density and with no helpers achieved higher within-pair paternity success and males that were in higher density areas gained more EPP. Additionally, all (three) cases of incestuous pairings resulted in EPP (though not all nestlings in these broods were EP), suggesting the possibility that females may partially avoid inbreeding depression by mating extra-pair when partnered with a close relative.

## CONCLUSION

We found support for plumage colouration to have a signalling function in mate choice in a dichromatic species where both sexes are ornamented. More colourful individuals invested more in parental care, and females and males paired and mated assortatively by plumage colour, suggesting mutual mate choice. Lovely fairy-wrens appear to have sex differences in their reproductive strategies and its relationship with plumage colouration: while no significant relationship was found between female plumage colour and EP offspring (male mate choice), less colourful males were more successful in getting extra-pair paternity (female mate choice). Our results are inconsistent with more conspicuous plumages being a signal of higher reproductive success. Instead, less colourful plumages in males seem to confer a reproductive advantage, at least in the context of extra-pair mating success. It is possible that females may be more selective than males and choose social and extra-pair males based on their plumage colour, or males adopt different mating strategies (EPP vs parental care) that are related with plumage colour, which ultimately have similar fitness. More empirical work is needed to understand how and what drives females and male mate choice for plumage colour.

Taking these results together with previous work (Leitão et al., 2019a), plumage colour in both sex seems to function in intra and inter-sexual competition contexts, although the information conveyed by plumage colouration seems to be contextdependent in relation to the sex of the bearer, which might partially explain dichromatism in this species. This study contributes to the growing body of the research on the function and evolution of female traits but stresses the need for further work comparing both female and male in a single system, so we can better understand sexdifferences in traits, possible mechanisms driving these differences, and their functional and evolutionary consequences.

## Acknowledgments

We acknowledge the Traditional Custodians of the land on which this work was conducted, the Yirrganydji people of the Djabugay country and the Wurundjeri people of the Kulin Nation. We would like to thank Rachel Shepperd, Gaia Marini, Phil Chaon, Irene Cuesta, Zoe Zelazny, Luke Nelson, and Kelsey Bell for field assistance over 2015-2017, and Brian Venables for field support. Thank you to Timon van Asten and Andrew Katsis for support in the laboratory during DNA extraction, Derick Thrasher for developing and providing the *M. lamberti* microsatellites.

Our study was approved by the University of Melbourne animal ethics committee (register 1613868.1) following guidelines under national and international legislation for animal use in research. Fieldwork was carried out under licence from Queensland Parks and Wildlife Service (WISP13237913), and birds were banded under Australian Bird and Bat Banding Scheme banding permits.

Authors’ contributions: A.V.L., M.L.H. and R.M. developed the idea, A.V.L. collected the field data, analysed, and wrote the manuscript with the contribution of all authors.

Data accessibility: Analyses reported here can be reproduced using the data provided by AVL.

## SUPPLEMENTARY DATA

**Table A1.** Description of microsatellite loci screened for posterior panel, based on dominant birds.

| Locus | N<br>samples<br>run | Nr<br>Alleles | Size<br>range | %<br>Successful<br>samples | Variability |
| --- | --- | --- | --- | --- | --- |
| Mcymu1<sup>1</sup> | 22 | 2 | - | 0 | - |
| Mcymu4<sup>1</sup> | 35 | 4 | 190-196 | 94.29 | 0.12 |
| Mcymu6<sup>1</sup> | 35 | 8 | 151-161 | 0.28 | 0.28 |
| Mcymu7<sup>1</sup> | 230 | 6 | 118-126 | 98.26 | 0.03 |
| Mcymu8<sup>1</sup> | 230 | 26 | 258-353 | 94.35 | 0.12 |
| Msp4<sup>2</sup> | 22 | 3 | 169-169 | 0.60 | 0.60 |
| Msp6<sup>2</sup> | 35 | 5 | 279-291 | 0.17 | 0.14 |
| Msp10<sup>2</sup> | 35 | 2 | 189-189 | 0.06 | 0.06 |
| Tmm6<sup>3</sup> | 230 | 63 | 316-469 | 93.48 | 0.29 |
| Smm1<sup>4</sup> | 208 | 9 | 257-273 | 79.33 | 0.05 |
| Smm2<sup>4</sup> | 35 | 4 | 208-218 | 0.11 | 0.11 |
| Smm7<sup>4</sup> | 35 | 3 | 262-264 | 0.09 | 0.09 |
| Ase9<sup>5</sup> | 13 | 4 | 143-149 | 0.31 | 0.31 |
| MaLa2<sup>6</sup> | 215 | 20 | 203-295 | 98.60 | 0.09 |
| MaLa3<sup>6</sup> | 20 | 3 | 297-301 | 0.15 | 0.15 |
| MaLa4<sup>6</sup> | 215 | 7 | 337-353 | 98.60 | 0.03 |
| MaLa5<sup>6</sup> | 215 | 8 | 193-217 | 97.67 | 0.04 |
| MaLa6<sup>6</sup> | 215 | 7 | 190-206 | 98.60 | 0.03 |
| MaLa7<sup>6</sup> | 215 | 7 | 290-310 | 98.14 | 0.03 |
| MaLa8<sup>6</sup> | 215 | 10 | 233-269 | 98.14 | 0.05 |
| MaLa10<sup>6</sup> | 215 | 11 | 271-326 | 98.14 | 0.08 |
| MaLa13<sup>6</sup> | 215 | 11 | 331-363 | 98.60 | 0.05 |
| MaLa14<sup>6</sup> | 215 | 11 | 250-282 | 98.14 | 0.05 |
| MaLa16<sup>6</sup> | 20 | 4 | 186-187 | 0.22 | 0.22 |
| MaLa18<sup>6</sup> | 215 | 8 | 258-290 | 98.60 | 0.04 |

References for microsatellite are: <sup>1</sup>(Double et al., 1997); <sup>2</sup>(Webster et al., 2004); <sup>3</sup>(Adcock and Mulder, 2002); <sup>4</sup>(Maguire et al., 2006); <sup>5</sup>(Richardson et al., 2000); <sup>6</sup>(Thrasher et al., 2017).

**Table A2.** Loadings for principal component analysis (PC1_chroma_) of xyz colour for cheek and head for each sex separately. Variable x represents the relative stimulation of S cone in relation to VS cone; y represents relative stimulation of M cone in relation to VS and S cones; and z axis represents the relative stimulation of L cone in relation to VS, S and M cones

| | Variables | $PC1_{\text{chroma}}$ | |
| --- | --- | --- | --- |
|  |  | Female | Male |
| **Cheek** | x | -0.05 | 0.12 |
|  | y | -0.25 | -0.68 |
|  | z | -0.96 | -0.71 |
|  | Variance | 91 % | 73 % |
| **Head** | x | -0.13 | -0.04 |
|  | y | -0.01 | -0.09 |
|  | z | -0.99 | -0.99 |
|  | Variance | 99 % | 95 % |

**Table A3.** Repeatability of colour measurement between years using the rptR package (Nakagawa and Schielzeth, 2010). Significant values in bold.

|  | *R* | SE | *CI* | *P* | *LRT* |
| --- | --- | --- | --- | --- | --- |
| Head Colour<br>N = 9 | 0.54 | 0.23 | [0-0.86] | **0.05** | -62.40 |
| Cheek colour<br>N = 11 | 0.36 | 0.22 | [0-0.76] | 0.13 | -59.50 |

**Table A4.** General linear model for body condition and body size. Significant predictors in bold. *N* = 34

|  | Female |  |  |  | Male |  |  |  |
| --- | --- | --- | --- | --- | --- | --- | --- | --- |
| | $\beta$ | SE | $t$ | $P$ | $\beta$ | SE | $t$ | $P$ |
| **Body condition** |  |  |  |  |  |  |  |  |
| Intercept | 8.59 | 0.20 | 42.50 | **<0.001** | 8.85 | 0.08 | 104.81 | **<0.001** |
| Cheek colour | -0.02 | 0.10 | -0.25 | 0.80 | -0.04 | 0.06 | -0.67 | 0.50 |
| Head colour | 0.03 | 0.05 | 0.66 | 0.51 | 0.01 | 0.03 | 0.27 | 0.78 |
| **Body size** |  |  |  |  |  |  |  |  |
| Intercept | 20.44 | 0.18 | 112.12 | **<0.001** | 21.35 | 0.13 | 161.14 | **<0.001** |
| Cheek colour | 0.12 | 0.09 | 1.33 | 0.19 | -0.13 | 0.10 | -1.24 | 0.223 |
| Head colour | -0.07 | 0.04 | -1.64 | 0.11 | 0.03 | 0.05 | 0.65 | 0.51 |

**Table A5.**
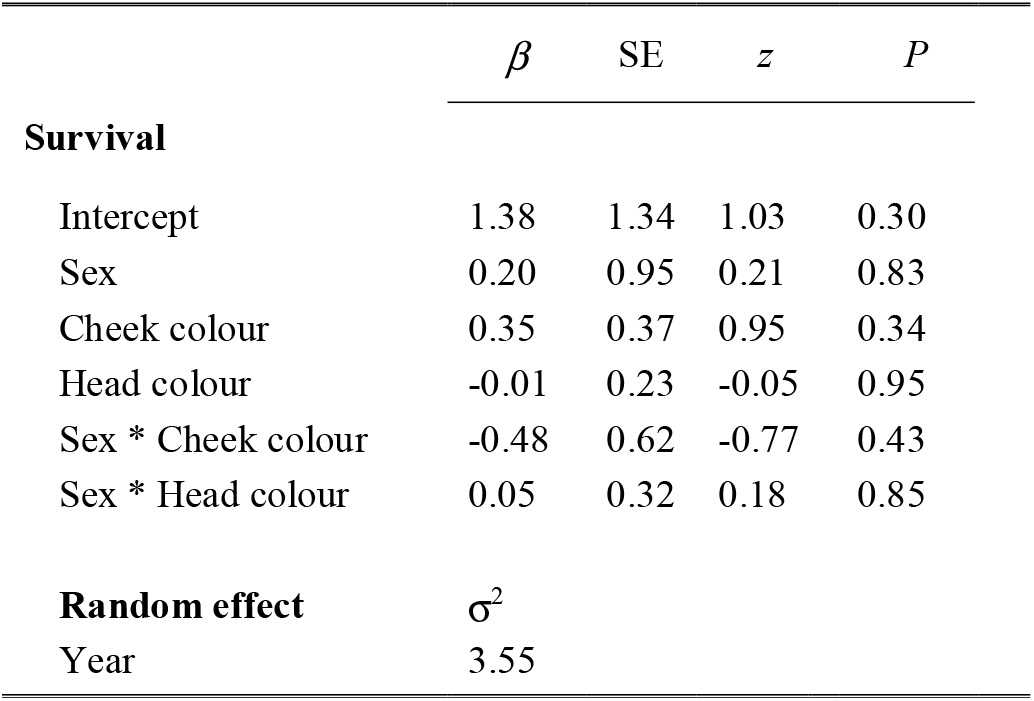
General linear mixed model for survival (survived more than 12 months after capture/colour measurement = 1; died before one year = 0). *N* = 74

